## Supplementary_File1 for "Exposing distinct subcortical components of the auditory brainstem response evoked by continuous naturalistic speech"

***Supplementary Table 1A.*** *LMER model formula and summary output for multiband peaky speech in experiment 1.*

Formula: log10(latency – a) ~ wave + log10(frequency) + wave:log10(frequency) + (1 + log10(frequency) | subject)

| Coefficient | Estimate | SEM | t | p | | Power (95% CI) |
|---|---|---|---|---|---|---|
| ***High-pass 30 Hz*** | | | | | | |
| (Intercept) | 0.596 | 0.012 | 51.5 | < 0.001 | *** | 1.00 (0.99, 1.00) |
| wave P0 | 0.516 | 0.011 | 45.1 | < 0.001 | *** | 1.00 (0.99, 1.00) |
| wave Na | 0.667 | 0.012 | 57.6 | < 0.001 | *** | 1.00 (0.99, 1.00) |
| wave Pa | 0.776 | 0.012 | 66.2 | < 0.001 | *** | 1.00 (0.99, 1.00) |
| log10(frequency) | -0.413 | 0.022 | -18.6 | < 0.001 | *** | 1.00 (0.99, 1.00) |
| wave P0:log10(frequency) | 0.189 | 0.027 | 7.0 | < 0.001 | *** | 1.00 (0.99, 1.00) |
| wave Na:log10(frequency) | 0.262 | 0.028 | 9.3 | < 0.001 | *** | 1.00 (0.99, 1.00) |
| wave Pa:log10(frequency) | 0.317 | 0.029 | 11.0 | < 0.001 | *** | 1.00 (0.99, 1.00) |
| ***High-pass 150 Hz*** | | | | | | |
| **Coefficient** | **Estimate** | **SEM** | **t** | **p** | | **Power (95% CI)** |
| (Intercept) | 0.597 | 0.012 | 50.9 | < 0.001 | *** | 1.00 (0.99, 1.00) |
| wave P0 | 0.487 | 0.012 | 40.7 | < 0.001 | *** | 1.00 (0.99, 1.00) |
| wave Na | 0.638 | 0.012 | 51.5 | < 0.001 | *** | 1.00 (0.99, 1.00) |
| wave Pa | 0.761 | 0.013 | 60.6 | < 0.001 | *** | 1.00 (0.99, 1.00) |
| log10(frequency) | -0.373 | 0.022 | -16.8 | < 0.001 | *** | 1.00 (0.99, 1.00) |
| wave P0:log10(frequency) | 0.137 | 0.030 | 4.6 | < 0.001 | *** | 0.99 (0.98, 1.00) |
| wave Na:log10(frequency) | 0.173 | 0.034 | 5.1 | < 0.001 | *** | 1.00 (0.99, 1.00) |
| wave Pa:log10(frequency) | 0.217 | 0.035 | 6.1 | < 0.001 | *** | 1.00 (0.99, 1.00) |

*p < 0.05, **p < 0.01, ***p < 0.001, CI = confidence interval

Note 1: (Intercept) and log10(frequency) are terms that refer to the default condition, which is wave V. Estimates of interactions of these terms reflect the change compared to the default. Frequency was the center frequency of the band normalized to 1 kHz (i.e., center frequency / 1000)

Note 2: a = τsynaptic + τI-V. Tsynaptic = 0.8 and τI-V was the subjects' I-V interval from their response to peaky broadband speech. If a subject did not have a wave I then the mean I-V was used for that subject.

***Supplementary Table 1B.*** *LMER model formula and summary output for multiband peaky speech in experiment 2.*

Formula: log10(latency – a) ~ wave + narrator + log10(frequency) + wave:narrator + wave:log10(frequency) + narrator:log10(frequency) + (1 + log10(frequency) | subject)

| ***High-pass 30 Hz*** | | | | | | |
|---|---|---|---|---|---|---|
| **Coefficient** | **Estimate** | **SEM** | **t** | **p** | | **Power (95% CI)** |
| (Intercept) | 0.646 | 0.018 | 35.4 | < 0.001 | *** | 1.00 (0.99, 1.00) |
| female narrator | -0.051 | 0.015 | -3.3 | 0.001 | *** | 0.93 (0.92, 0.95) |
| wave P0 | 0.472 | 0.017 | 27.6 | < 0.001 | *** | 1.00 (0.99, 1.00) |
| wave Na | 0.607 | 0.018 | 31.7 | < 0.001 | *** | 1.00 (0.99, 1.00) |
| wave Pa | 0.733 | 0.018 | 41.0 | < 0.001 | *** | 1.00 (0.99, 1.00) |
| log10(frequency) | -0.488 | 0.028 | -15.9 | < 0.001 | *** | 1.00 (0.99, 1.00) |
| female narrator:log10(frequency) | 0.008 | 0.024 | 0.3 | 0.732 | | 0.06 (0.05, 0.08) |
| wave P0:log10(frequency) | 0.201 | 0.032 | 6.2 | < 0.001 | *** | 1.00 (0.99, 1.00) |
| wave Na:log10(frequency) | 0.271 | 0.033 | 8.1 | < 0.001 | *** | 1.00 (0.99, 1.00) |
| wave Pa:log10(frequency) | 0.300 | 0.034 | 8.8 | < 0.001 | *** | 1.00 (0.99, 1.00) |
| wave P0:female narrator | 0.035 | 0.021 | 1.7 | 0.087 | | 0.42 (0.39, 0.45) |
| wave Na:female narrator | 0.020 | 0.021 | 0.9 | 0.346 | | 0.18 (0.16, 0.20) |
| wave Pa:female narrator | 0.009 | 0.021 | 0.4 | 0.685 | | 0.07 (0.06, 0.09) |

*p < 0.05, **p < 0.01, ***p < 0.001, CI = confidence interval

Note 1: (Intercept) and log10(frequency) are terms that refer to the default condition, which is wave V for the male narrator. Estimates of interactions of these terms reflect the change compared to the default. Frequency was the center frequency of the band normalized to 1 kHz (i.e., center frequency / 1000)

Note 2: a = τsynaptic + τI-V. Tsynaptic = 0.8 and τI-V was the subjects' I-V interval from their response to peaky broadband speech. If a subject did not have a wave I then the mean I-V was used for that subject.

***Supplementary Table 1C.*** *LMER model formula and summary output for multiband peaky speech in experiment 3.*

Formula: log10(latency – a) ~ narrator + log10(frequency) + narrator:log10(frequency) + (1 + log10(frequency) | subject)

| ***High-pass 30 Hz*** | | | | | | |
|---|---|---|---|---|---|---|
| **Coefficient** | **Estimate** | **SEM** | **t** | **p** | | **Power (95% CI)** |
| (Intercept) | 0.615 | 0.015 | 40.0 | < 0.001 | *** | 1.00 (0.99, 1.00) |
| right ear | 0.013 | 0.012 | 1.1 | 0.284 | | 0.21 (0.18, 0.23) |
| female narrator | -0.041 | 0.014 | -2.9 | 0.004 | *** | 0.83 (0.80, 0.85) |
| log10(frequency) | -0.360 | 0.020 | -18.4 | < 0.001 | *** | 1.00 (0.99, 1.00) |
| female narrator:log10(frequency) | -0.047 | 0.027 | -1.7 | 0.085 | | 0.40 (0.37, 0.43) |

*p < 0.05, **p < 0.01, ***p < 0.001, CI = confidence interval

Note 1: (Intercept) and log10(frequency) are terms that refer to the default condition, which is wave V for the male narrator and left ear. Estimates of interactions of these terms reflect the change compared to the default. Frequency was the center frequency of the band normalized to 1 kHz (i.e., center frequency / 1000)

Note 2: a = τsynaptic + τI-V. Tsynaptic = 0.8 and τI-V was the mean I-V interval from the response to peaky broadband speech in experiment 2.
